# SUITOR: selecting the number of mutational signatures through cross-validation

**DOI:** 10.1101/2021.07.28.454269

**Authors:** Donghyuk Lee, Difei Wang, Xiaohong R. Yang, Jianxin Shi, Maria Teresa Landi, Bin Zhu

## Abstract

For *de novo* mutational signature analysis, the critical first step is to decide how many signatures should be expected in a cancer genomics study. An incorrect number could mislead downstream analyses. Here we present SUITOR (Selecting the nUmber of mutatIonal signaTures thrOugh cRoss-validation), an unsupervised cross-validation method that requires little assumptions and no numerical approximations to select the optimal number of signatures without overfitting the data. *In vitro* studies and *in silico* simulations demonstrated that SUITOR can correctly identify signatures, some of which were missed by other widely used methods. Applied to 2,540 whole-genome sequenced tumors across 22 cancer types, SUITOR selected signatures with the smallest prediction errors and almost all signatures of breast cancer selected by SUITOR were validated in an independent breast cancer study. SUITOR is a powerful tool to select the optimal number of mutational signatures, facilitating downstream analyses with etiological or therapeutic importance.

## Background

Mutational signatures are patterns of somatic mutations imprinted on the cancer genome by operative mutational processes. At least 49 single base substitution mutational signatures have been identified across cancer types^1^, with some associated with exogenous mutagenic exposures^2-4^ and endogenous mutational processes^5-8^. Moreover, mutational signatures have been proposed to identify cancer predisposition genes (e.g., *NTHL1* in breast cancer^9^), and to stratify cancer patients^10-13^ for precision treatment. In these studies, deciding the expected number of signatures is the pivotal first step, which determines the downstream steps of extracting signature profiles, estimating signature contributions and stratifying tumors based on signatures for treatment. As an example, the Pan-Cancer Analysis of Whole Genomes (PCAWG) consortium reported that the discordance between the extracted and known signatures is usually caused by the difficulty in selecting the correct number of signatures^1^.

Despite the many algorithms proposed to extract signature profiles^14-18^ and estimate signature contributions^19,20^, relatively little emphasis has been placed on selecting the correct number of *de novo* mutational signatures in cancer genomics studies^21,22^. SomaticSignatures^14^ chooses the number of signatures based on the residual sum of squares and the explained variance without a clearly defined selection criterion. SigProfiler^15^ considers the mean reconstruction error and the stability of signature extraction; however, it is unclear how these features could be combined to jointly predict the number of signatures. EMu^16^ and signeR^17^ adopt a Bayesian information criterion (BIC)^23^. Although BIC is a popular model selection criterion for supervised learning (e.g., regression and classification) where the number of parameters is fixed, it may not be applicable to unsupervised learning, including mutational signature analysis, where the number of parameters increases with the sample size (other limitations of BIC elaborated in Supplementary Note 1). SignatureAnalyzer^1^ uses an automatic relevance determination (ARD) prior^24^ which imposes a sparsity assumption on mutation profiles and contributions. The number of signatures chosen by signatureAnalyzer is sensitive to the pre-specified sparsity assumption, especially hyperparameters of the ARD prior and the tolerance level.

To overcome the limitations of previous methods, we propose selecting the number of signatures through cross-validation. Selecting the number of signatures is essentially a problem of model selection, which has been addressed by cross-validation in other research areas^25,26^, including identification of cancer subtypes^27^, exploration of population structure^28^ and prediction of lymph node metastasis^29^. In the setting of mutational signature analysis, cross-validation splits the full dataset (here, the mutation counts) into a training set and a validation set; for a given number of signatures, these signatures are estimated in the training set and then they are used to predict the mutations in the validation set. Multiple candidate numbers of signatures are considered; and the number of signatures which predicts most closely the mutations in the validation (not the training) set is selected. Hence, cross-validation can prevent selecting too few or too many signatures (corresponding to an underfitting or overfitting model), both of which would predict mutation counts in the validation set poorly. In addition, unlike the BIC or the ARD prior, cross-validation requires little assumptions and no numerical approximations^26^. Therefore, cross-validation provides a viable solution for selecting the correct number of signatures.

Despite being conceptually appealing, the standard cross-validation approach does not work for unsupervised mutational signature analysis. In the standard cross-validation scheme, it is feasible to remove a subset of subjects all together as a validation set because parameters (e.g., regression coefficients) are identical across subjects. However, mutation contributions in mutational signature analysis are tumor-specific parameters. If mutation counts of a subset of tumors were removed all together as a validation set, these counts would be used twice (for estimating tumor-specific mutation contributions and calculating the prediction error), which would underestimate the prediction error. Instead, cross-validation for mutational signature analysis requires retaining all tumors in the training set but removing some mutation counts from each tumor as a validation set. Consequently, missing data emerge in the training set, which would cause current methods for mutational signature analysis to fail.

These limitations are overcome by SUITOR (Selecting the nUmber of mutatIonal signaTures thrOugh cRoss-validation), an unsupervised cross-validation method that selects the optimal number of signatures to attain the minimal prediction error in the validation set. SUITOR extends the probabilistic model to allow missing data in the training set, which makes cross-validation feasible. Moreover, we propose an expectation/conditional maximization (ECM) algorithm^30^ to extract signature profiles, estimate mutation contributions and impute the missing data simultaneously. We demonstrated SUITOR’s superior performance using *in vitro* experimental data, *in silico* simulations, *in vivo* applications to 2,540 tumors across 22 cancer types, and validation of signatures of breast cancer in additional 440 breast tumors.

## Results

### Overview of SUITOR

Mutational signature analysis decomposes the catalog matrix of somatic mutations. Take single base substitution (SBS) as an example. A mutation catalog matrix **V** of size N × 96 contains mutation counts for N cancer genomes and 96 SBS catalogs. Each SBS catalog refers to a mutated pyrimidine (C or T) in the center and two unmutated adjacent nucleotides (flanking 5’ and 3’ bases) with total 4 × 6 × 4 = 96 catalogs. For example, a genomic sequence A<u>C</u>G in the normal tissue is mutated to A<u>G</u>G in the tumor tissue. This SBS belongs to the A[C > G]G mutation catalog.

SUITOR is built upon a probabilistic model^31,32^, for which the maximum likelihood estimation (MLE) is equivalent to the solution of non-negative matrix factorization (NMF), the most popular method for mutational signature analysis^15^. Given the number of signatures *r* to be extracted, NMF factorizes the mutation catalog matrix into two non-negative matrices: signature contribution matrix **W** of size N × *r* and signature profile matrix **H** of size *r* × 96 such that **V ≈ WH**. Each row of **W** contains attributions of *r* signatures, reflecting how intense *r* mutational signatures are in a tumor; each row of **H** forms a signature profile with the elements summed to 1, showing how 96 mutation catalogs comprise a signature profile. To estimate **W** and **H**, it is common to minimize the generalized Kullback-Leibler (KL) divergence

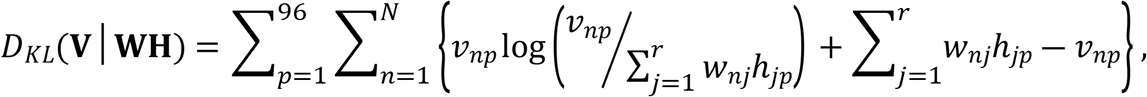

subject to *w*_*nj*_ ≥ 0 and *h*_*jp*_ ≥ 0 with 1 ≤ *j* ≤ r, 1 ≤ *n* ≤ N and 1 ≤ *p* ≤ 96. Lowercase letters, *v*_*np*_, *w*_*nj*_ and *h*_*jp*_, denote elements of the corresponding matrices, **V, W** and **H**, respectively. NMF can also be solved with other objective functions such as Frobenius norm or more general *β*-divergence, depending on the applications^24^.

Notably, minimization of the generalized KL divergence is equivalent to maximize a likelihood function of a probabilistic NMF model^32,33^. Indeed, for a Poisson NMF model, *v*_*np*_ of the *n*th tumor and *p*th mutation catalog is assumed to be independently distributed, following a Poisson distribution with mean 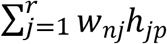. The log-likelihood of Poisson NMF model can be written as log {Pr(**V**|**WH**)} = − *D*_*KL*_(**V**|**WH**) +*C* with a constant *C*. Therefore, minimizing generalized KL divergence *D*_*KL*_(**V**|**WH**) is equivalent to maximizing the log-likelihood log {Pr(**V**|**WH**)}. In addition, it could been shown that the multiplicative update algorithm^34^, which is commonly used to minimize the generalized KL divergence, is equivalent to an expectation/conditional maximization (ECM) algorithm^30^ for the Poisson NMF model (Supplementary Note 2). These two equivalences are used to develop SUITOR.

As mentioned above, NMF or equivalently the Poisson NMF model requires a given number of signatures *r*, which is unknown in practice; and SUITOR can select this number empirically. The steps of SUITOR are outlined as follows (with a schematic illustration in Fig.1): 1) the mutation catalog matrix is separated into the training and validation sets. The training set contains missing data held out as validation data; 2) the missing data are imputed in the initial step; 3) the ECM algorithm iteratively imputes the missing data in the expectation step (E-step) and estimates the signature contributions (of the **W** matrix) and profiles (of the **H** matrix) in the conditional maximization steps (CM-steps) until the ECM algorithm allows **V ≈ WH** for a given number of signatures; 4) the missing data are imputed and compared to the validation data to calculate the prediction error in the validation set; 5) The above described steps are conducted for multiple candidate numbers of signatures and the one with the minimal prediction error will be chosen as the optimal number of signatures (corresponding to the red dot in the prediction error curve of validation set in Fig. 1).

**Fig.1.**
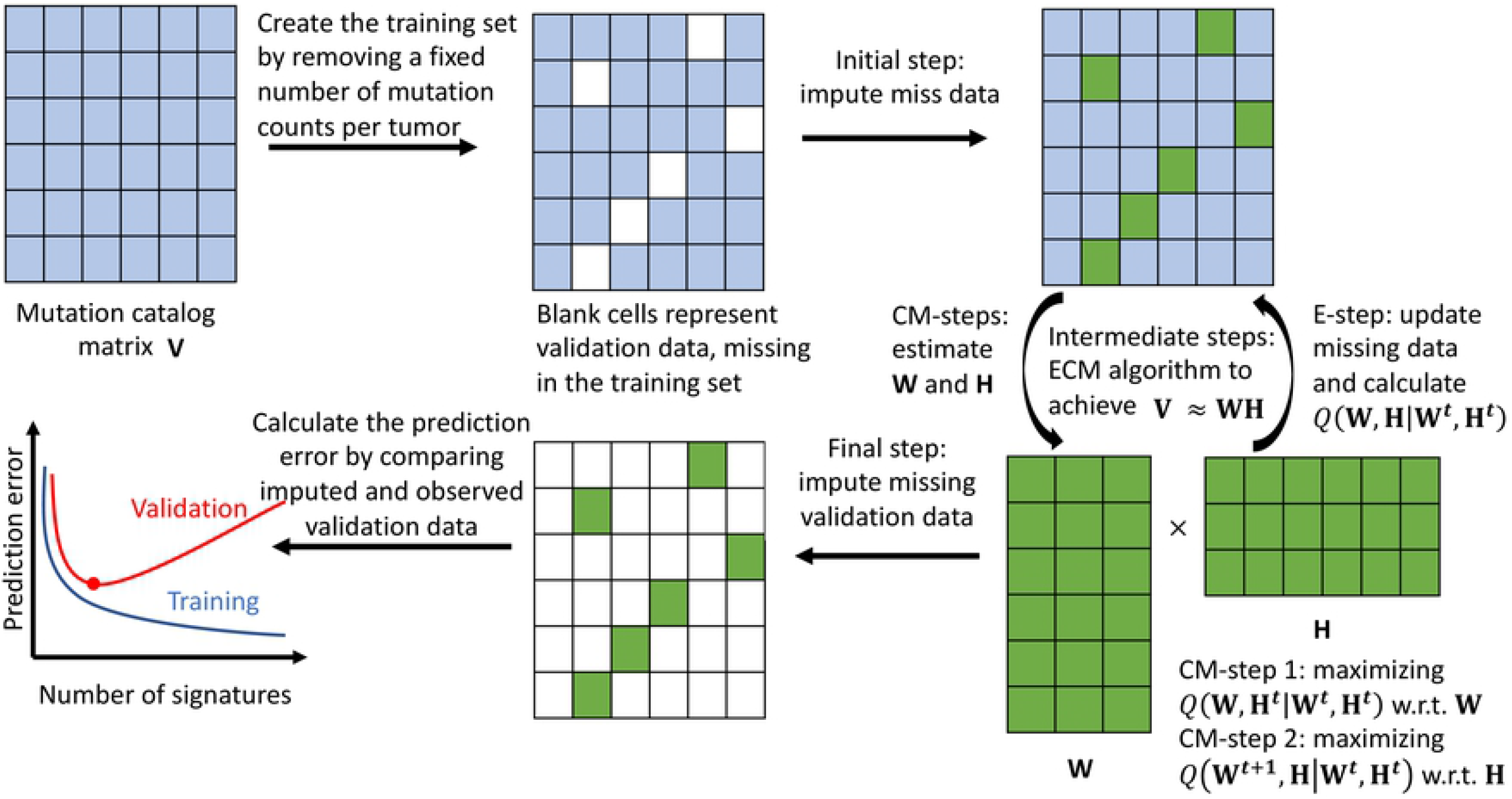
A schematic overview of SUITOR. This schematic diagram illustrates how SUITOR selects the number of *de novo* mutational signatures. Details are given in Results. Each row of the mutation catalog matrix ***V*** represents a tumor, each column a mutation catalog. The bottom left shows prediction error curves in the training set (blue) and validation set (red), which are manually drawn for the illustration purpose, with the red dot representing the minimal prediction error in the validation set. ECM algorithm: expectation/conditional maximization algorithm; CM-steps: conditional maximization steps; E-step: expectation step; **W**: signature contribution matrix of size N × *r* , N the number of tumor and *r* the number of signatures; **H**: signature profile matrix of size *r* × 96; *Q*(***W***,***H***|***W***^***t***^,***H***^***t***^): conditional expectation of complete likelihood of **W** and **H** at the (*t* + 1)-th step, given estimated ***W***^***t***^ and ***H***^***t***^ in the *t*-th step of ECM algorithm.

There are three key contributions of SUITOR. First, SUITOR selects the number of signatures with the minimal prediction error, which prevents selecting too few or too many signatures (as model underfitting or overfitting). Although the prediction error in the training set is reduced with increasing signatures (as illustrated by the prediction error curve of the training set in Fig. 1), the prediction error in the validation set will decrease first (due to the model underfitting with insufficient signatures) and then inflate (due to the model overfitting with redundant signatures). This is the well-known bias-variance tradeoff for model complexity^35^ measured by the number of signatures in the setting of mutational signature analysis. Second, the cross-validation scheme of SUITOR guarantees that the missing data pattern does not depend on the remaining or missing mutation counts in the training set. Hence, the missing data mechanism is missing completely at random (MCAR), which ensures that the estimated signature profiles and contributions would not be biased due to the missing data^36^. Third, the proposed ECM algorithm for SUITOR enjoys the convergence property, which guarantees the increase of the likelihood function over iterations until the ECM algorithm converges^30^.

### Evaluation of SUITOR in two *in vitro* studies

We assessed the performance of SUITOR in two experimental studies^2,5^, for which the true number and profile of signatures were generated experimentally and validated *in vitro* (see details in Methods). The first study created endogenous mutational signatures through CRISPR-Cas9-mediated knockouts of DNA repair genes in an isogenic human cell line^5^. The second study generated exogenous mutational signatures in human-induced pluripotent stem cell (iPSC) lines exposed to environmental or therapeutic mutagens^2^. For both studies, we evaluated whether SUITOR could correctly select the number of signatures and recover the profiles of single base substitution signatures. We then compared SUITOR’s performance with sigProfiler, signatureAnalyzer and signeR.

For the *in vitro* CRISPR-Cas9-mediated knockout study of the DNA repair gene *MSH6*, SUITOR correctly detected the background signature, which existed before the knockout of *MSH6* and *MSH6* knockout-induced signature (Fig. 2a) and recovered the corresponding signature profiles (Fig. 2b; Supplementary Table 1). SigProfiler, signatureAnalyzer and signeR correctly identified these two signatures as well (Supplementary Fig. 1; Supplementary Table 1). Next, we extracted signatures of knockout studies of six DNA repair genes (*CHEK2, NEIL1, NUDT1, POLB, POLE* and *POLM)*, which did not induce experimentally detectable signatures^5^. SUITOR and the other three methods correctly identified one background signature only without false detection of knockout-induced signatures (Supplementary Fig. 2). We conclude that in this *in vitro* study with at most two signatures, all four methods perform equally well. This is not the case when the number of signatures increases as shown below.

**Fig.2.**
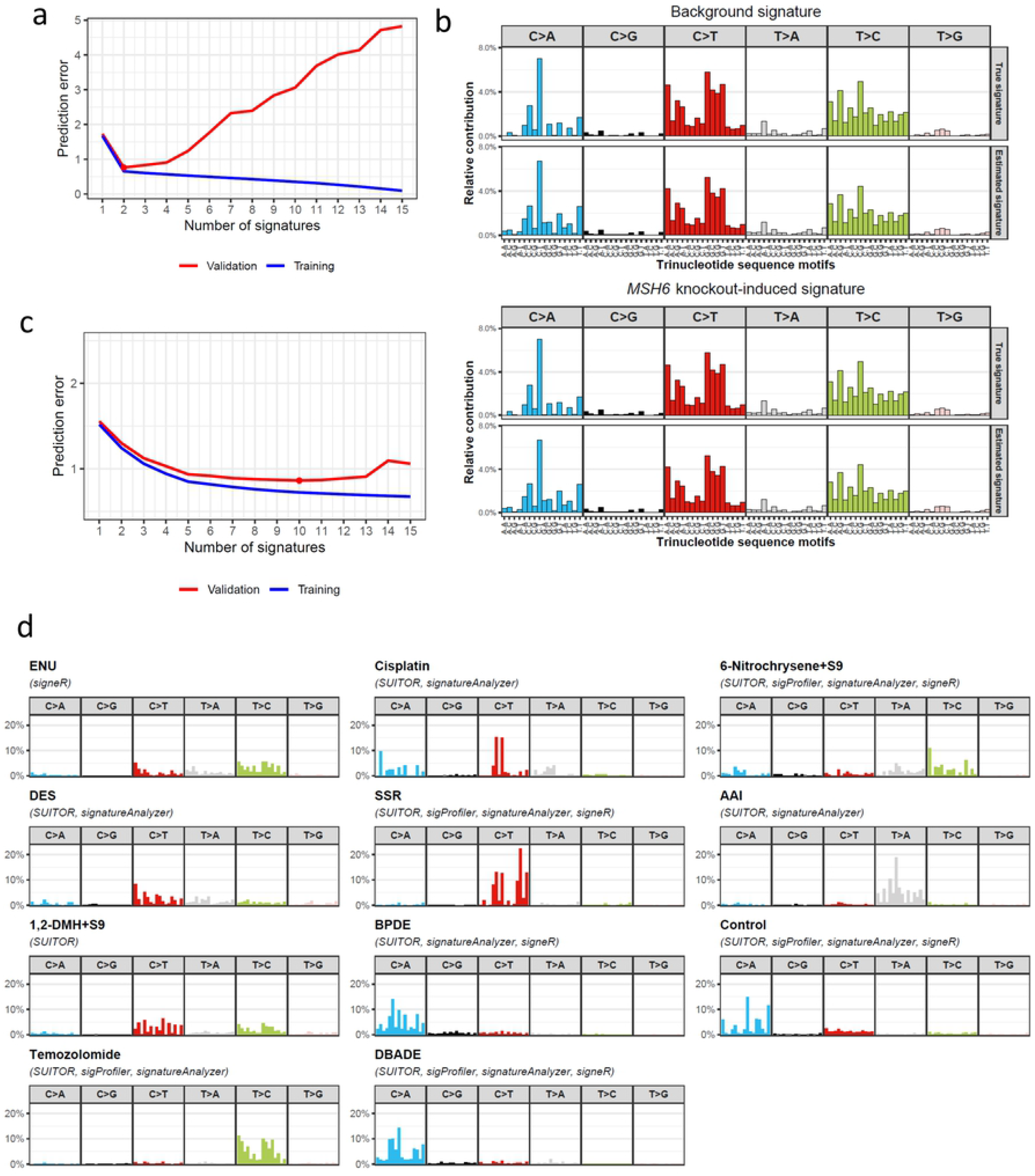
*In vitro* evaluation of SUITOR and other methods. a) Prediction errors of SUITOR for the training and validation sets of *in vitro* knockout study of DNA repair gene *MSH6*. The red dot denotes the number of signatures with the minimal prediction error in the validation set. b) Profiles of single base substitution signature estimated by SUITOR in *MSH6* gene knockout study. The x-axis indicates the 5’ and 3’ nucleotides for each substitution type (e.g., T[C>A]C, cytosine to adenine substitution with 5’ thymine and 3’ cytosine). The top panel: the true and estimated background signatures (cosine similarity=0.991; cosine similarity=1 suggests two profiles being identical.); the bottom panel: the true and estimated *MSH6* knockout-induced signatures (cosine similarity=0.997). c) Prediction errors by SUITOR for the training and validation sets of *in vitro* study of environmental or therapeutic mutagens. The red dot indicates the minimal prediction error in the validation set achieved by ten signatures, including one background signature and nine mutagen-induced signatures. d) Signatures discovered from the *in vitro* study of environmental or therapeutic mutagens by four methods. Methods which could identify a given mutagen are enclosed in paratheses. ENU: N-ethyl-N-nitrosourea; 6-Nitrochrysene+S9: 6-Nitrochrysene mixed with S9 rodent liver-derived metabolic enzyme; DES: diethyl sulfate; SSR: simulated solar radiation; AAI: aristolochic acid I; 1,2-DMH+S9: 1,2-dimethylhydrazine mixed with S9 rodent liver-derived metabolic enzyme; BPDE: benzo[a]pyrene-7,8-dihydrodiol-9,10-epoxide; DBADE: dibenz[a,h]anthracene diol-epoxide.

For the *in vitro* study of 79 exogenous mutagens, stable mutational signatures were experimentally identified for 28 mutagens. These 28 signatures are not distinct as some signatures are very similar to each other (e.g., benzo[a]pyrene (BaP) and benzo[a]pyrene-7,8-dihydrodiol-9,10-epoxide (BPDE), both of which are polycyclic aromatic hydrocarbons (PAHs), mutagens of tobacco smoke; Supplementary Fig. 3). Although all four methods could find the background mutational signature, SUITOR detected 9 additional signatures induced by mutagens (Figure 2c, 10 signatures, including one background signature and 9 mutagen-induced signatures), sigProfiler detected 4, signatureAnalyzer 8, and signeR 5 additional signatures (Figure 2d; Supplementary Fig. 4; Supplementary Table 2); the signature of 1,2-dimethylhydrazine (1,2-DMH) was detected by SUITOR and missed by the other methods. Besides detecting more true signatures, SUITOR also achieved the lowest prediction error (SUITOR:1292.5; sigProfiler:2295.4; signatureAnalyzer:2176.6; signeR:1585.1). This indicates that the signatures detected by SUITOR in the training set predicted most closely the mutation counts in the validation set. When we stratified the tumor subclones based on the signatures detected by at least one method, SUITOR and signatureAnalyzer were able to separate most subclones into distinct clusters, each corresponding to a unique mutagen exposure (Supplementary Fig. 5). In contrast, SigProfiler merged subclones exposed to BPDE and dibenz[a,h]anthracene diol-epoxide (DBADE); signeR mixed subclones exposed to N-ethyl-N-nitrosourea (ENU) and 1,2-DMH; and both SigProfiler and signeR mixed subclones exposed to 9-Nitrochrysense and aristolochic acid I (AAI).

We note that *in vitro* study of exogenous mutagens favors the method of signatureAnalyzer. SignatureAnalyzer implicitly assumes few signatures are present per sample and hence the loadings of signatures are sparse, which holds here since each sample was treated with a single mutagen in this *in vitro* study of exogenous mutagens. In spite of that, SUITOR performed better than signatureAnalyzer. When the sparsity assumption does not hold, as demonstrated in *in silico* simulations and the PCAWG study described below, signatureAnalyzer would find more false-positive signatures than the other methods. In contrast, SUITOR, which does not rely on the sparsity assumption, is not susceptible to it.

### *In silico* simulation studies

We evaluated SUITOR and three other methods through additional *in silico* simulations. First, we evaluated the scenario when only random mutations occur. SUITOR identified one random signature in 20 of 20 replicates while other methods found many more signatures (as many as 96 signatures for signatureAnalyzer; Supplementary Table 3). Note that the random signature identified by SUITOR is similar to SBS3 (cosine similarity = 0.87), implying that SBS3 would possibly be caused by random mutations. Next, we considered a setting of one true signature, for which any additional signature found would be a false positive (details in Methods). SUITOR, sigProfiler and signeR correctly identified the single true signature (Supplementary Fig. 6) in 20 of 20 replicates; signatureAnalyzer found a false positive signature (Fig. 3a) in 17 of 20 replicates. Finally, we examined a setting of nine true signatures with varying signature contributions (details in Methods; Supplementary Fig. 7; Supplementary Table 4). All methods correctly identified six common signatures (SBS1, 2, 3, 5, 13, 18) in 20 of 20 replicates; SUITOR and signatureAnalyzer detected two rare signatures (SBS8,41) in more replicates than the other two methods (Fig. 3b). None of the methods identified the extremely rare and flat signature SBS40 (present in one tumor only). Notably, all methods detected no other signatures besides the nine true signatures. Together, the simulation studies suggest that SUITOR is able to find both common and rare signatures while well controlling the rate of false positives.

**Fig.3.**
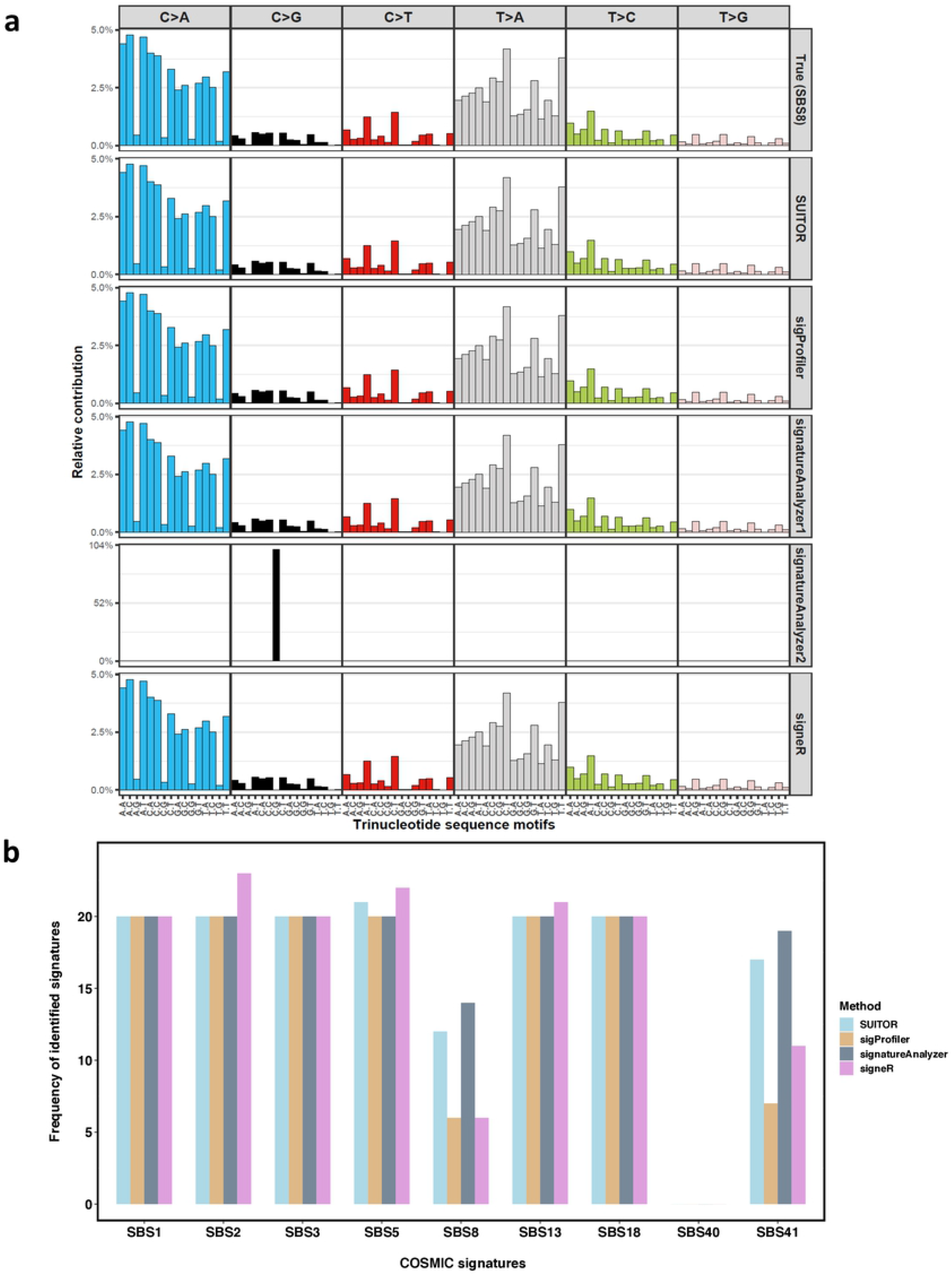
*In silico* evaluation of SUITOR and other methods. a) The signature profiles of the true mutational signature and the ones discovered by each method for a replicate. b) The number of replicates in which a given signature will be detected by each method; the rare signatures SBS8 and SBS41 were discovered in 12, 6, 14 and 6 replicates, and 17, 7, 19 and 11 replicates over total 20 replicates by SUITOR, sigProfiler, signatureAnalyzer and signeR, respectively.

### Detection of pan-cancer mutational signatures

We tested the four methods in whole-genome sequencing (WGS) data of 2,540 tumors across 22 cancer types from the Pan-Cancer Analysis of Whole Genomes (PCAWG) study^1^ (details in Methods).

First, we extracted *de novo* mutation signatures one cancer type at a time for eight cancer types, each with at least 100 tumors. Unlike in *in vitro* or *in silico* studies, the true signatures were unknown here. Nevertheless, we could evaluate if the signatures detected in part of the dataset predict mutation counts in the remaining part well. Specifically, the mutation catalog matrix is separated into training, validation and testing sets, the last of which is used to evaluate the performance of the selected number of signatures (details in Methods). Among the four methods, SUITOR clearly attained the smallest prediction errors across eight cancer types (Fig. 4a). Moreover, most signatures found by SUITOR were highly similar to the COSMIC signatures (with cosine similarity > 0.8, Fig. 4b) and frequently detected by the other methods (Fig. 4c). In contrast, signatureAnalyzer identified more *de novo* signatures, some of which were not matched to any COSMIC signatures (Fig. 4b).

**Fig.4.**
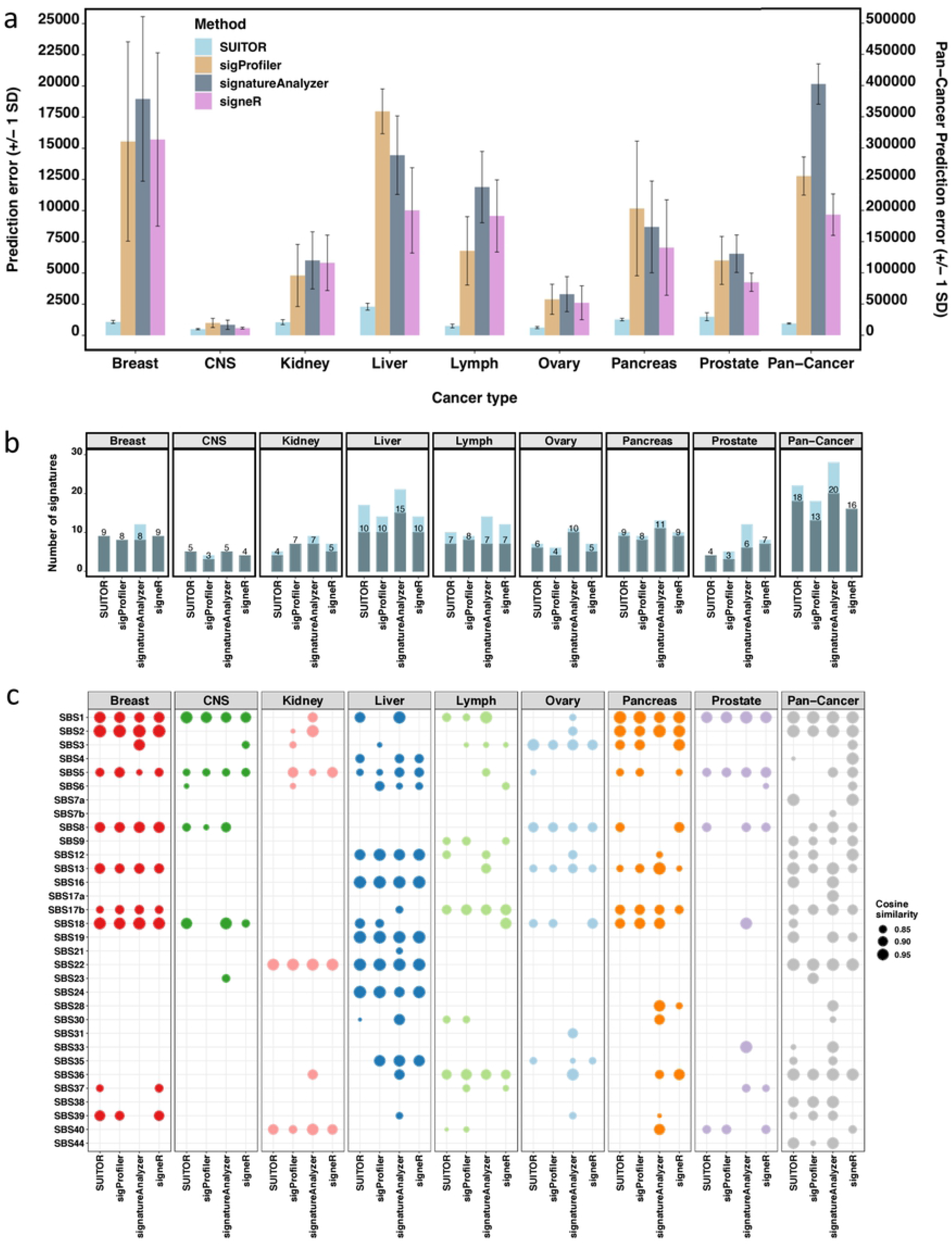
Mutational signature results of eight cancer studies of PCAWG. a) The prediction errors of SUITOR, sigProfiler, signatureAnalyzer and signeR for eight cancer types (scale on the left-side Y axis) and for 2,540 tumors across 22 cancer types together, namely Pan-Cancer (scale on the right-side Y axis). SD: standard deviation. b) The number of signatures identified by each method. The shaded bars and the numbers above indicate the number of signatures whose profiles could be matched to COSMIC profiles (with cosine similarity > 0.8). c) The cosine similarities between *de novo* signatures and COSMIC signatures. Only the matched pairs are shown (with cosine similarity > 0.8). The higher the cosine similarity the better match to a COSMIC signature profile. The cosine similarity equivalent to one denotes a perfect match.

Next, we extracted *de novo* signatures combining WGS data from 2,540 tumors (pan-cancer analysis). SUITOR found 22 signatures, eighteen of which could be matched to the COSMIC signatures (Fig. 4b). These signatures had the smallest prediction error (Fig. 4a), compared to signatures detected by the other methods: SUITOR 36,017; sigProfiler 255,111 (>7 times SUITOR’s prediction error); signatureAnalyzer 402,409 (>11 times SUITOR’s prediction error); and signeR 193,191 (>5 times SUITOR’s prediction error). As expected, the signatures commonly found in multiple cancer types (e.g., SBS1, SBS2 and SBS13) could be identified when combining all cancer types together, while signatures specific to a single cancer type (e.g., SBS24 specific to liver cancer) were absent in the combined signature analysis (Fig. 4c). When we clustered 2,540 tumors based on signature contributions estimated by SUITOR, four clusters emerged in the t-SNE plot (Fig. 5a). Liver tumors formed a distinct cluster, possibly due to its specific signature SBS24 caused by aflatoxin exposure; the remaining clusters included: i) a subset of lymphomas; ii) the majority of kidney tumors; iii) all remaining tumors. Although the liver tumors were separated from the other tumors in the t-SNE plots also by the other methods (Fig. 5b-d), the separation of lymphomas and kidney tumors was less clear. For example, kidney tumors and lymphoma were mixed with other cancer types in the t-SNE plot from signatureAnalyzer.

**Fig.5.**
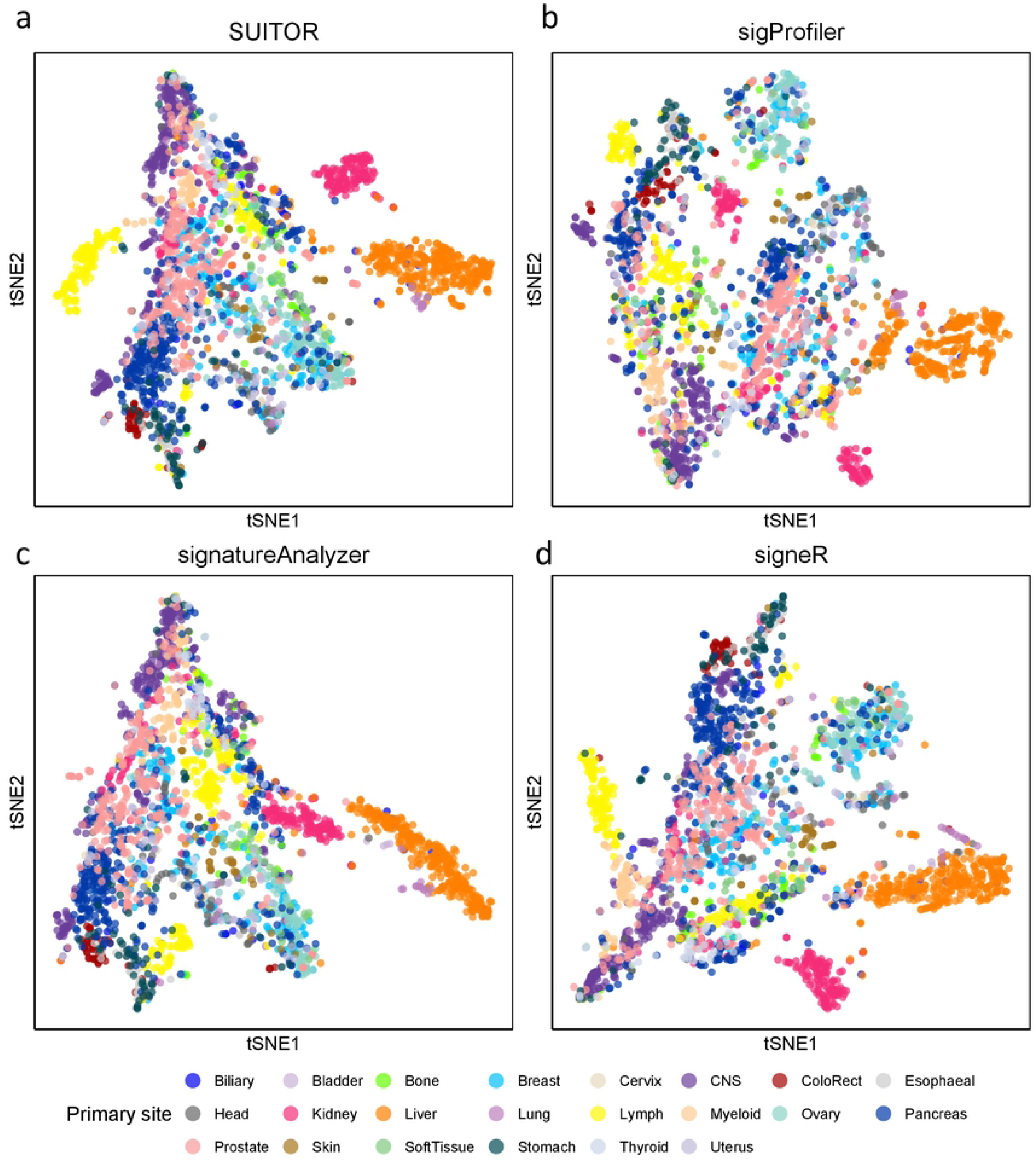
The t-SNE visualization of clustering patterns for 2,540 tumors across 22 cancer types based on signature contributions. Signature contributions are estimated by four methods: (a) SUITOR, (b) sigProfiler, (c) signatureAnalyzer and (d) signeR. Each dot represents a tumor and is colored by the cancer type.

### External validation of breast cancer mutational signatures

We validated the nine PCAWG breast cancer signatures (based on 194 breast tumors) using an independent WGS set of 440 breast tumors of the Sanger breast cancer (BRCA) study from the same ethnicity^37^. SUITOR (Fig. 6a and Supplementary Table 5) and signeR (Supplementary Table 6) identified nine breast cancer signatures in PCAWG and validated eight in the Sanger BRCA study (with cosine similarity > 0.8); sigProfiler identified eight signatures and confirmed seven (Supplementary Table 7); signatureAnalyzer identified twelve and validated eight (Supplementary Table 8). Overall, six signatures (SBS1,2,8,13,17b and 18) were identified in both studies by all methods, while the flat featureless signature SBS5 could not be validated by any method.

**Fig.6.**
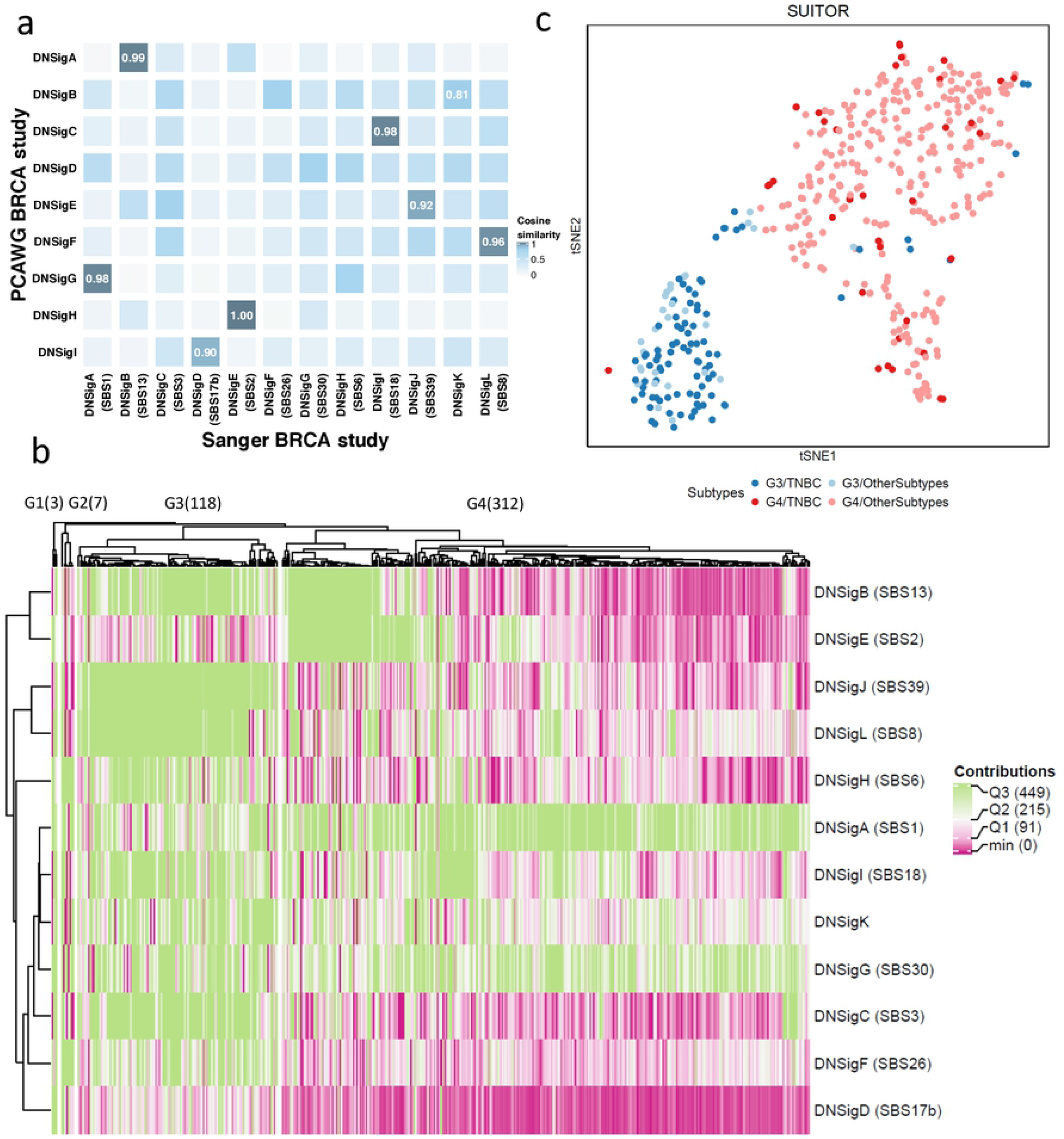
The results of Sanger breast cancer study by SUITOR. a) The heatmap of cosine similarity between *de novo* signatures detected in the PCAWG breast cancer (BRCA) study and *de novo* signatures in the Sanger BRCA study. The signatures of the Sanger BRCA study are annotated by COSMIC signatures (if cosine similarities > 0.8) among parentheses. For example, DNSigA (SBS1) refers to *de novo* signature A (DNSigA) being annotated by COSMIC signature SBS1. b) The heatmap of signature contributions estimated by SUITOR with hierarchical clustering in the Sanger BRCA study. The number of tumors included in each signature cluster is shown among parentheses on the top of the heatmap. Q1, Q2, Q3: the 1st, 2nd and 3rd quantiles of signature contributions. c) The t-SNE visualization of tumors in G3 and G4 signature clusters, color-coded by signature clusters and molecular subtypes. TNBC: triple negative breast cancer.

Besides validating eight PCAWG breast cancer signatures, SUITOR found four additional signatures in the Sanger BRCA study (Fig. 6a and Supplementary Table 9). Two of them were highly similar to COSMIC signatures SBS26 and SBS30 (cosine similarity > 0.93) and were identified by the other three methods as well; the other two (similar to SBS3 and SBS6) were also detected by signatureAnalyzer and/or signeR. Similarly, three other methods found few more signatures as well (Supplementary Tables 10-12). These findings suggest that there likely exist additional signatures in the Sanger BRCA study that are missed in the PCAWG breast cancer study because of either the larger sample size or specific operative mutational processes (e.g., BRCA1 mutation carriers with signature SBS3) in Sanger BRCA study.

Finally, as an example of clinical utility of the signatures estimated by SUITOR, we stratified the 440 breast tumors of the Sanger BRCA study using the signature contributions. Four signature clusters were found; two dominant clusters (G3 and G4) included overall 430 tumors (Fig. 6b). Compared to the G4 subgroup, the G3 subgroup showed significantly higher contributions of the nine signatures (Supplementary Fig. 8; Supplementary Table 13) and significantly lower contributions of the de novo signature A (similar to COSMIC signature SBS1 associated with aging). We found a number of clinical factors associated with the subgroups G3 and G4, including age at diagnosis and tumor grade (Supplementary Table 14). The singular most important associated factor was the molecular subtype: triple negative breast cancers were significantly enriched in the subgroup G3 (Fig. 6c; odds ratio = 25.1, P-value < 2.2×10^−16^, two-sided Fisher’s exact test).

## Discussion

It’s crucial to select the correct number of *de novo* mutational signatures for a cancer genomics study. Here we present SUITOR that selects the number of signatures through cross-validation to minimize the prediction error in the validation set. We have shown how SUITOR outperforms common existing methods most of the time. *In vitro* studies show that SUITOR is capable of retrieving the correct number and profiles of both endogenous and exogenous signatures, allowing the correct stratification of tumor subclones exposed to distinct mutagens. *In silico* simulation studies show that SUITOR can detect common signatures in all replicates and rare signatures (as low as 1%) in the majority of replicates. Applications to *in vivo* eight PCAWG cancer types show that SUITOR discovers signatures which achieve the lowest prediction errors in the testing sets. Most of these signatures are confirmed by other methods and matched to the COSMIC signatures. All except one signature found in PCAWG BRCA study were validated in the independent Sanger BRCA study. The contributions of signatures selected by SUITOR in the Sanger BRCA study are dominated by two clusters, driven by the molecular/histological subtypes.

Besides selecting the number of signatures of single base substitution, SUITOR could be used to select the number of signatures of other genomic alterations in tumors, including double base substitutions, small insertion and deletions (indels), and structure variations. In this paper, we used 10-fold cross validation, which is recommended as a good compromise for the bias-variance trade-off regarding the choice of k in k-fold cross validation^38,39^. In addition, we have tried 20-fold cross validation (i.e., 5% of data as validation data) for PCAWG studies, which led to the same number of signatures (results not shown).

In summary, SUITOR has shown to perform better than other commonly used methods in revealing mutational signatures, the “footprints” engraved in the cancer genomes by operative mutational processes with potentially important etiological or therapeutic implications.

## Methods

### Unsupervised cross-validation for mutational signature analysis

SUITOR aims to select the optimal number of signatures which minimizes the prediction error in the validation set through cross-validation. Here, we describe the steps to create the validation set and the related challenges.

For a *K*-fold cross-validation, we divide the catalog matrix **V** into K parts where the Poisson NMF model is fitted on *K* −1 parts as the training set, and the fitted model is validated on the remaining one part as the validation set. The cross-validation is carried out K times with each part served as a validation set once, using the balanced separation^40,41^ detailed as follows. In the *k*^*th*^ fold (1 ≤ *k* ≤ *K*) of a balanced separation, a set of mutation counts {*v*_*np*_|*p* = (*n mod* 10) + (*k* − 1) + *aK,a* = 1, 2, ….} are held out for the *n*^*th*^ tumor as validation data, where *a* is restricted such that 1 ≤ *p* ≤ 96. For example, (1, 11, …, 91)^*st*^ mutation catalogs of the first tumor are held out in the 1^*st*^ fold, (2, 12, …, 92)^*nd*^ mutation catalogs in the 2^*nd*^ fold and so on. Note that the balanced separation keeps equal number of retained mutation catalogs for each tumor in the training set, which is computationally more stable than randomly splitting **V** into the training and validation sets. The latter may randomly remove a large number of mutation catalogs for a tumor.

As validation data are hold out, missing data emerge in the training set, the reason for which existing methods of NMF fail. To address this challenge, we extended the Poisson NMF model and propose an expectation/conditional maximization (ECM) algorithm to incorporate the missing data.

### Expectation/conditional maximization (ECM) algorithm of SUITOR

Let 𝒮 be the set of indices of mutation catalog matrix **V** such that 𝒮 = {(*n,p*)|1 ≤ *n* ≤ N and 1 ≤ *p* ≤ 96}. For a *K*-fold cross-validation, 𝒮 would be divided into *K* disjoint sets 𝒮_1_, …,𝒮_*K*_. For the *k*^*th*^ fold, 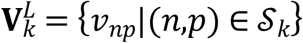 denotes the validation set and 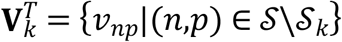 the training set, where 𝒮\𝒮_*k*_ represents the indices of **V** excluding ones in 𝒮_*k*_. Mutation counts in 𝒮_*k*_ will be removed from **V** and denote as missing data **M**_*k*_ = {*m*_*np*_|(*n,p*) ∈ 𝒮_*k*_}. The *v*_*np*_ in 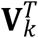 and *mnp* in **M**_*k*_ are assumed to be independently distributed as a Poisson distribution with the mean 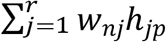. By the scheme of balanced separation, 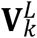 is missing completely at random (MCAR), since 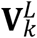 is removed from **V**, independent of values of 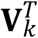 and 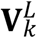. MCAR enables us to propose an ECM algorithm to incorporate missing data and obtain unbiased estimates of **W** and **H** (ref36).

Next, we outline the ECM algorithm in the following iterative steps (details in Supplementary Note 3).

1. Initial step: choose initial values of **M**_*k*_ and set initial parameters **W**^**0**^ and **H**^**0**^.
2. E-step: given the observed data 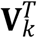 and the parameters **W**^***t***^ and **H**^***t***^ of the previous step *t*, the ECM algorithm calculates the conditional expectation of complete likelihood

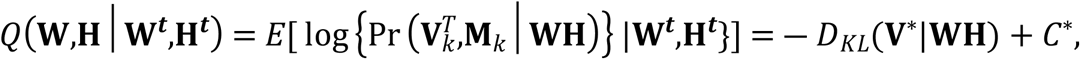

where *C*^*^ is a constant independent of **W** and **H**, 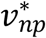 the elements of **V**^*^ as 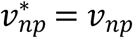 for (*n,p*) ∈ 𝒮/𝒮_*k*_ and 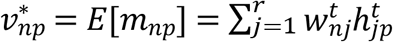 for (*n,p*) ∈ 𝒮_*k*_
3. CM1-step: update parameters **W**^***t+1***^by maximizing *Q*(**W**,**H**^***t***^|**W**^***t***^,**H**^***t***^) with respect to **W**.
4. CM2-step: update parameters **H**^***t+1***^by maximizing *Q*(**W**^***t***+**1**^,**H**|**W**^***t***^,**H**^***t***^) with respect to **H**.
5. Iterate steps 2 to 4 until convergence.

In the initial step, we use the median of mutation counts in 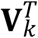 per each mutation catalog as initial values for **M**_*k*_; other more complicated methods of specifying initial values, such as nearest neighbors, lead to similar results (results not shown). We are aware that the ECM algorithm possibly converges to a local saddle point. To overcome it, we try 300 random initial values **W**^**0**^ and **H**^**0**^, which leads to 300 pairs of ***Ŵ***_*i*_ and ***Ĥ***_*i*_, the estimates of **W** and **H** for the *i*th initial value, *i* = 1, 2,…, 300. The final reported **Ŵ** and **Ĥ** are the ***Ŵ***_*i*_ and ***Ĥ***_*i*_ which maximize the 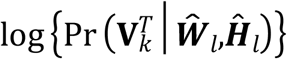 among all 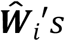 and 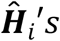.

### Selecting number of signatures by SUITOR

For a given number of signatures *r*, we first evaluate the prediction error, i.e., the disparity between the observed validation data 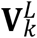 and the predicted ones 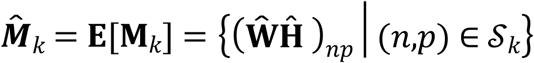, for the *k*^*th*^ fold, *k* = 1,2,…,*K*

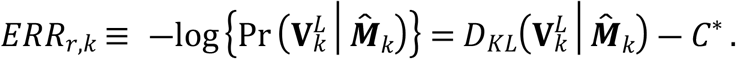

We then evaluate overall prediction error, 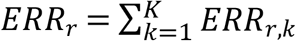, across K folds. Since the term

*C*^*^ is unrelated to **Ŵ** and **Ĥ**, it is dropped. Finally, we select the number of signatures *r*^*^ which minimizes *ERR*_*r*_ over a range of numbers of signatures 1 ≤ *r* ≤ *R*.

### Extracting signature profiles and estimating contributions of signatures

Once the optimal number of signatures *r*^*^ is determined by SUITOR, we extract mutational signature profiles **W** and estimating contributions **H**, via maximizing log {Pr(**V**|**WH**)} with the fixed rank *r*^*^. Similar to the ECM algorithm in SUITOR, we evaluate multiple initial values and use ***Ŵ*** and ***Ĥ*** which maximizes log {Pr (**V Ŵ** |**Ĥ)}**to relieve local optima problem.

### *In vitro* studies

The datasets of two *in vitro* studies were downloaded from http://medgen.medschl.cam.ac.uk/serena-nik-zainal/. The details of study design and implementation were described previously^2,5^. In these *in vitro* studies, the endogenous and exogenous mutational signatures were experimentally generated *in vitro* and hence the true number of signatures and profiles are known.

We created the mutation catalog matrix for both studies and applied SUITOR, sigProfiler, signatureAnalyzer and signeR. We chose sigProfiler and signatureAnalyzer, since they have been applied to a number of studies^7,8,18,42-44^, and signeR^17^ because it utilizes Bayesian information criterion (BIC), a popular model selection criterion for supervised learning. For SUITOR, we used 10-fold cross-validation with 90% of counts in mutation catalog matrix as the training set and the remaining 10% as the validation set. In contrast, the whole mutation catalog matrix **V** was analyzed by sigProfiler, signatureAnalyzer and signeR, respectively under the default setting.

The first study induced endogenous mutational signatures by CRISPR-Cas9-mediated knockouts of DNA repair genes in an isogenic human cell line. First, we focused on the *MSH6* knockout-induced single base substitution signature, which is characterized by C>T and T>C single base substitutions (∼148 substitutions per cell division). We evaluated whether the four methods were able to retrieve the background signature and the *MSH6* knockout-induced signatures. Next, we analyzed the gene-knockout studies with no induced signatures (for genes *CHK2, NEIL1, NUDT1, POLB, POLE* and *POLM*), to evaluate whether the four methods would find false positive signatures in addition to the background signature.

In the second study, exogenous mutational signatures were created by environmental or therapeutic mutagens. We selected 324 subclones (including 35 control subclones) of human-induced pluripotent stem cell (iPSC) lines, for which the mutations were measured by whole-genome sequencing (WGS). While controls are not exposed to mutagens, each subclone is exposed to one of 79 mutagens, including simulated solar radiation (SSR), dibenzo[a,l]pyrene (DBP) and alkylating agent therapy temozolomide (TMZ). SSR recapitulates the UV-associated signatures and DBP is a potent carcinogen of the polycyclic aromatic hydrocarbons (PAHs) produced when coal, crude oil, or gasoline is burned. For each method, we checked if the correct number of signatures were attained with its impacts on the downstream analyses. Specifically, we investigated whether the retrieved *de novo* signature profiles were highly similar to the true signature profiles. We further explored if signature contributes could separate subclones exposed to the distinctive mutagens, visualized by t-distributed stochastic neighbor embedding (t-SNE^45^).

### *In silico* simulation design with random mutations

We simulated mutation catalog matrix for 300 tumors directly from the Poisson distribution with the mean lambda, which in turn was drawn from a uniform distribution and varied from 20 to 160. We repeated this simulation 20 times and analyzed the simulated catalog matrices by SUITOR, sigProfiler, signatureAnalyzer and signeR.

### *In silico* simulation design with one signature

We simulated a mutation catalog matrix for 500 tumors and analyzed it by the four methods, each repeated 20 times. We used the signature profile of SBS8 as the true signature profile for **H** and generated the contribution vector **W** from a uniform distribution within the range [20000, 40000]. Then the mutation catalog **V** was generated by a Poisson distribution with mean **WH**.

### *In silico* simulation design with nine signatures

We simulated signatures mimic to the ones observed in the Pan-Cancer Analysis of Whole Genomes (PCAWG) breast cancer study^1^. The nine signatures identified by SUITOR show various signature contributions and signature profiles; some signatures contribute to all tumors (e.g., SBS1 and SBS5, present in 100% of tumors; Supplementary Table 4) while others contribute to a few tumors (e.g., SBS41, present in 6% of tumors) and even to one (SBS40) or two tumors (SBS8); some signature profiles are spiky (e.g., SBS1 and SBS2/13; Supplementary Fig. 5) while others are relatively flat (e.g., SBS5).

Specifically, we generated mutation catalog matrices similar to **V**^BR^, the mutation catalog matrix of PCAWG breast cancer study. **V**^BR^ is approximated by **W**^BR^**H**, for which **H** contains the COSMIC signature profiles and **W**^BR^ is the corresponding signature contribution matrix (downloaded from https://www.synapse.org/#!Synapse:syn11738669). We removed signatures with zero contributions to all tumors and chose 9 signatures to compose the signature profile matrix **H** of size 9 × 96. With **H** fixed, we took bootstrap samples 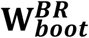 from each column of matrix **W**^BR^, and generated 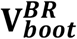 which follows a Poisson distribution with mean 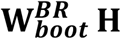. Due to the dependencies between SBS1 and SBS5 as well as between SBS2 and SBS13, their contributions are resampled jointly while the contributions of other signatures are sampled individually. We simulated 20 mutation catalog matrices for 200 tumors, and each was analyzed by four methods respectively.

### *In vivo* human cancer genomics studies

We analyzed 2, 540 tumors across 22 cancer types of PCAWG^1^, including 321 tumors of hepatocellular carcinoma, 286 tumors of prostate adenocarcinoma, 237 tumors of pancreatic adenocarcinoma, 194 tumors of breast adenocarcinoma, 146 tumors of central nervous system medulloblastoma, 143 tumors of renal cell carcinoma,112 tumors of ovary adenocarcinoma and 100 cases of B-cell non-Hodgkin lymphoma. Other cancer types have less than 100 tumors per cancer type. The tumors were whole genome sequenced and datasets of the somatic mutation calls were downloaded from https://www.synapse.org/#!Synapse:syn11726620. The hypermutator tumors with mutation burden more than 10 mutations/Mb were excluded^46^. For each cancer type, we applied SUITOR, sigProfiler, signatureAnalyzer and signeR to select the number of signatures and estimate signature contributions and profiles.

To compare the prediction errors, we split the mutation catalog matrix into a training set (90% of counts in the mutation catalog matrix), a validation set (5%) and a testing set (5%). For SUITOR, the training set was used to fit the probabilistic NMF model with multiple numbers of signatures and the validation set to select the number of signatures. The other methods used both the training and validation sets to select the number of signatures. Next, we compared the prediction errors of selected signatures by each method on the testing set. For sigProfiler, signatureAnalyzer and signeR, we imputed missing training data by medians of available mutation counts per each mutational catalog, applied each method, predicted the testing data and calculated the prediction error as SUITOR did. For SUITOR, it could handle missing data and predict the testing data by the ECM algorithm.

### Sanger whole genome sequencing breast cancer study

The Sanger whole genome breast cancer (BRCA) study sequenced 560 breast tumors. The somatic mutation calls files were downloaded from https://medgen.medschl.cam.ac.uk/serena-nik-zainal/. Among 560 breast tumors, 110 tumors were included in PCAWG and hence excluded from this validation study. Ten hypermutator tumors were also excluded. We applied SUITOR, sigProfiler, signatureAnalyzer and signeR to a) select the number of signatures and estimate signature contributions and profiles; b) compare the signatures with ones detected in the PCAWG breast cancer study; and c) investigate if additional signatures are found in Sanger whole genome breast cancer study. In addition, we stratified the tumors based on mutation contributions and associated the signature clusters with epidemiological or clinical characteristics.

### Software webpages

SUITOR: https://github.com/binzhulab/SUITOR

sigProfiler: v1.0.6 is installed and used by following instructions in https://github.com/AlexandrovLab/SigProfilerExtractor.

signatureAnalyzer: version information is not available but is installed by following instructions in https://github.com/broadinstitute/getzlab-SignatureAnalyzer.

signeR: v1.12.0 is downloaded from https://bioconductor.org/packages/release/bioc/html/signeR.html.

sigProfiler and signatureAnalyzer are used in python version 3.7.5 while SUITOR and signeR are used in R version 3.6.3. Signature profiles plot is drawn based on R package MutationalPatterns from https://bioconductor.org/packages/release/bioc/html/MutationalPatterns.html with slight modification to add graphical parameters.

## Acknowledgment

This research was supported by the Intramural Research Program of the National Institutes of Health, National Cancer Institute, Division of Cancer Epidemiology and Genetics (DCEG). This study utilized the high-performance computational capabilities of the Biowulf Linux cluster at the National Institutes of Health, Bethesda, MD: https://biowulf.nih.gov. We would like to thank Bill Wheeler (Information Management Services) for computation support and Dr. Paul Albert (Biostatistics Branch of DCEG) and Dr. Ludmil Alexandrov (University of California San Diego) for helpful comments. Conflict of Interest: None declared.

